# Protein kinase F regulates the virulence of *Mycobacterium tuberculosis*

**DOI:** 10.64898/2025.12.04.692467

**Authors:** Flor Torres-Juarez, Shivangi Rastogi, David Young, Paul J. Baker, Viplov K. Biswas, Shannon E. Khan, D. Branch Moody, Katrin D. Mayer-Barber, Volker Briken

**Affiliations:** Inflammation and Innate Immunity Section, Laboratory of Clinical Immunology and Microbiology, National Institute of Allergy and Infectious Diseases, Bethesda, MD, USA; Department of Cell Biology and Molecular Genetics, University of Maryland, College Park, MD, USA; Division of Rheumatology, Immunity and Inflammation, Brigham and Women’s Hospital, Harvard Medical School, Boston, MA, USA; Centre for Innate Immunity and Infectious Diseases, Hudson Institute of Medical Research, Clayton, Victoria 3168, Australia and Monash University, Clayton, Victoria 3800, Australia

## Abstract

The serine/threonine protein kinase F (PknF) of *Mycobacterium tuberculosis* (Mtb) has poorly defined targets and functions but is involved in limiting NLRP3 inflammasome activation in murine macrophages and dendritic cells *in vitro*. The importance of PknF for the virulence of Mtb *in vivo* is not known. Here, we demonstrate that the Mtb CDC1551 deletion mutant of *pknF (*Δ*pknF*) expresses significantly increased levels of the lipid pthiocerol dimycoserosate (PDIM), a polyketide lipid with pro-virulence properties. The Δ*pknF* mutant strain, when compared to the Mtb and complemented strains, had a 100-fold increase in growth at day 28 and about a 10-fold increase in growth at days 90-98 in the lungs of mice. The increase in pulmonary bacterial loads after infection with Δ*pknF* strain was conserved even in *Nlrp3*-deficient mice, arguing that PknF modulates Mtb virulence independently of NLRP3-inflammasome activation in mice. Staining of lung sections revealed increased inflammation in the lungs of Δ*pknF* strain-infected mice when compared to Mtb and the complemented mutant strain. Highly susceptible B6.Sst1^S^ mice displayed decreased host resistance with significantly decreased survival when infected with the Δ*pknF* strain compared to the complemented strain or Mtb. In conclusion, our data suggest that expression of PknF, as a modulator of multiple downstream effector proteins, restricts the virulence of Mtb in the lungs of mice through an NLRP3 inflammasome-independent mechanism but potentially via suppressing expression of the virulence lipid, PDIM.

**Author summary:** The human pathogen *Mycobacterium tuberculosis* (Mtb) encodes 11 serine/threonine protein kinases (Pkn A-I,K and L). *In vitro*, PknF inhibits activation of innate immune responses in macrophages and dendritic cells. The importance of that protein for the virulence of the bacteria in the context of a live animal infection is unknown. We used a hypersusceptible mouse strain to analyze the impact of deleting the *pknF* gene on the virulence of the bacteria by monitoring the survival of the mice. We show that the Δ*pknF* deletion mutant kills mice faster than wild-type bacteria or Δ*pknF* mutant bacteria expressing a wild-type copy of the *pknF* gene (Δ*pknF*-C). Next, we analyzed the growth of the different bacterial strains in the infected mice and demonstrated that at day 28 and day 90-98 timepoints the Δ*pknF* mutant bacterial strains showed a significant increase in bacterial burden in the lungs, bronchoalveolar lavage fluid and the spleen. An analysis of the total lipids of the bacterial stocks showed that the Δ*pknF* Mtb bacteria increased expression of the virulence lipid, pthiocerol dimycoserosate (PDIM), suggesting a negative regulation of this lipid by PknF likely affecting bacterial virulence in the lungs of mice.

## Introduction

Modulation of innate immune activation after infection of host cells is a key virulence mechanism employed by intracellular bacterial pathogens, especially infection with *Mycobacterium tuberculosis* (Mtb). Here we study the role of the Mtb serine/threonine protein kinase F (PknF), based on *in vitro* data showing its role in the regulation of the NLRP3 inflammasome activation, which tightly regulates the production of cleaved, bioactive IL-1β (1, 2). IL-1β is an important innate cytokine critical for host resistance to Mtb (3, 4). Mice deficient in the expression of either IL-1α, IL-1β or the IL-1β receptor (IL1R1) are hyper susceptible to Mtb infections (3, 5–10). IL-1R1 confers host resistance in mice via *trans*-protection of infected cells by both immune and non-immune cells and not via infected cell-autonomous anti-microbial mechanisms (11). Although the precise mechanisms remain unclear*, in vivo*, a major function of IL-1β seems to be to induce host protective eicosanoids and to inhibit detrimental type I interferon responses to promote cooperative tissue immunity (5, 10). Nucleotide-binding and oligomerization domain-like receptors (NLRs) survey the host cell cytosol for the presence of pathogens, and different NLRs recognize various pathogen-derived components or host cell stress to then initiate formation of the inflammasome complex (12, 13). For example, NLRP3 is activated by intracellular stress, and notably, the efflux of potassium ions (K^+^) is a major trigger (14, 15). The NLRP3 inflammasome in human and murine macrophages is slightly activated during *in vitro* infections with Mtb (16–19) via a mechanism that involves NLRP11 and caspase-4/5 in human cells (20). *In vivo*, using the mouse model, the inflammasome complex is not involved in generating IL-1β in response to Mtb infections (3, 16, 17). This inflammasome-independent IL-1β production in the mouse could potentially be due to the capacity of Mtb to inhibit the NLRP3 and AIM2 inflammasome activation (21–24). Indeed, an Mtb deletion mutant of the *ptpB* gene, which is involved in inflammasome inhibition, showed an increase in IL-1β response in the lungs of infected mice and was attenuated for growth (24).

A genetic screen revealed that *pknF* is important for inhibition of the NLRP3 inflammasome in murine macrophages (1) and dendritic cells (2) during Mtb *in vitro* infections. PknF is one of 11 members of a family of serine/threonine kinases in Mtb (26, 27). These protein kinases serve broader regulatory roles in Mtb by phosphorylating downstream target proteins that have shared function, so that the parallel phosphorylation of multiple effector programs can globally regulate Mtb physiology (26–28). The 476 amino acid (aa) PknF protein has one predicted transmembrane domain and a cytosolic 260 aa domain that has high homology with serine/threonine kinase domains of eukaryotic proteins (27). Global O-phosphorylation analysis after deletion of *pknF* documented reduced phosphorylation on 17 sites, suggesting that they are direct targets of PknF (28). *In vitro* assays show that PknF may phosphorylate Rv1747, Rv0020C, KasB, HupB and GroEL1 (29–32). *Rv1747* is in an operon with *pknF* (*rv1746*), and the phosphorylation of the T150 and T208 residues of Rv1747 activates this presumptive ATP-binding cassette (ABC) transporter (33). Rv1747 is negatively regulated by Rv2623 (34) which is phosphorylated by PknF (28). Interestingly, the deletion of *rv1747* produces an attenuation phenotype *in vivo* (33, 35), whereas the deletion of the negative regulator *rv2623* results in hypervirulence (36). There is limited information on the importance of PknF for the virulence of Mtb *in vivo* (37). In the present study, we investigated how deletion of *pknF* affects the virulence of the bacteria in both susceptible and resistant mouse strains and specifically tested for connections with NLRP3 inflammasome functions based on our previous *in vitro* findings.

## Results

The continuous culture of Mtb in liquid medium often results in the loss of pthiocerol dimycocerosate (PDIM) (38, 39). PDIM is an important driver of Mtb virulence in the mouse model (40–42). Consequently, we analyzed PDIM levels in our CDC1551 wild-type Mtb, Δ*pknF* mutant, and complemented mutant Δ*pknF*-C strains by performing an extraction of total lipids and separating multiple PDIM molecular species on a thin-layer chromatography plate (Fig. 1A). Interestingly, the Δ*pknF* strain showed increased PDIM expression when compared to the wild-type strain and, importantly, the complemented mutant strain Δ*pknF*-C (Fig. 1A). These results demonstrated a complementable phenotype of increased PDIM biosynthesis in the *pknF* deletion mutant.

**Fig. 1:**
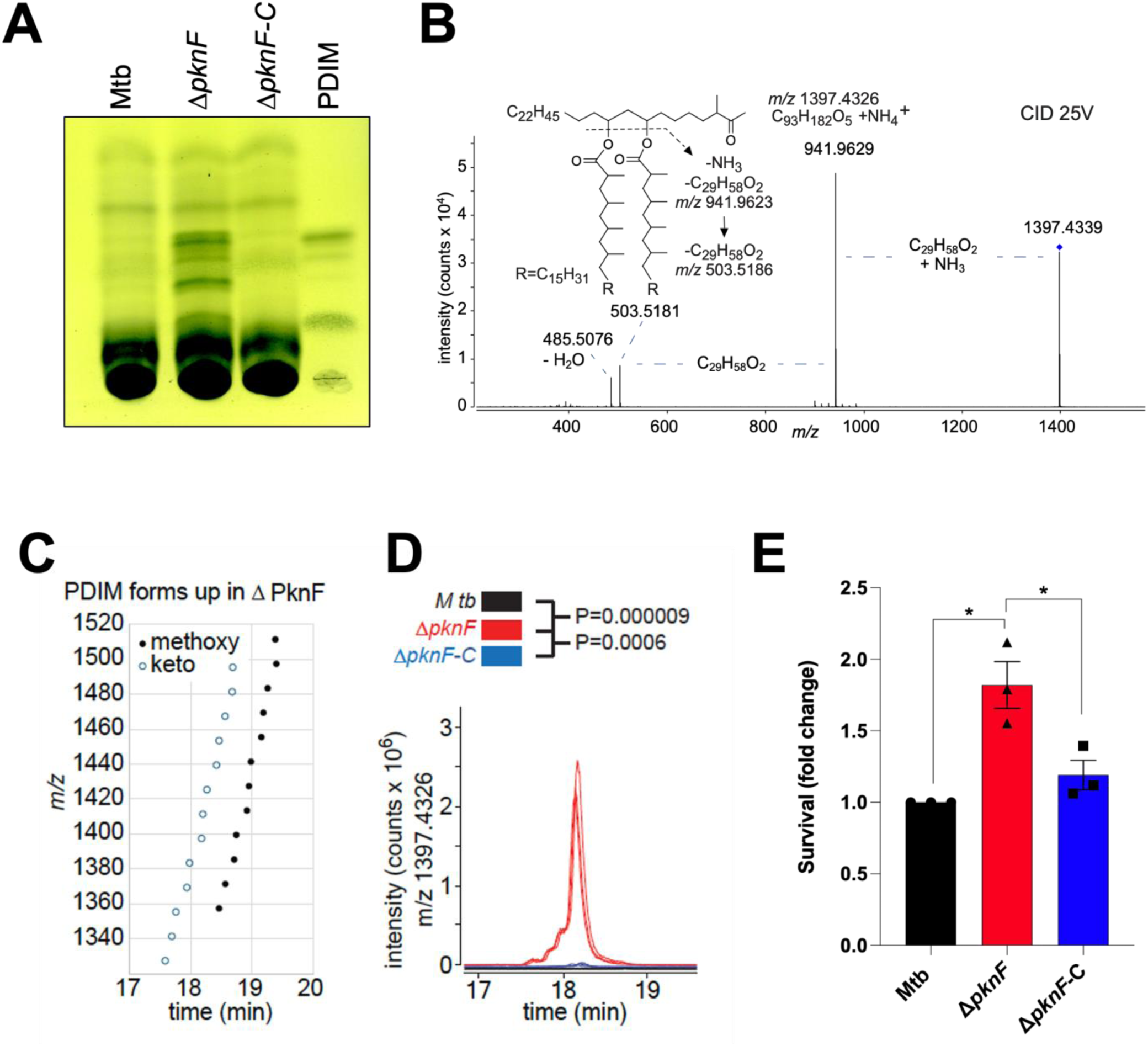
The deletion of *pknF* in Mtb CDC1551 causes an increase in PDIM expression. **(A)** Total lipids extracted from the indicated wild-type Mtb (CDC1551), the *pknF* deletion mutant (Δ*pknF*) and complemented mutant strains (Δ*pknF*-C) along with purified PDIM were separated by thin-layer chromatography. A representative result of three independent lipid extractions is shown. (**B**) The structure of PDIM, which is deduced at the keto form along with two C29 mycocerosic acid moities based on ESI-CID MS of ammonium adducts in the positive mode. (**C**) Fulfilling the expected biological distributions with chain length analogs that appear in alkane series deduced methoxy- and keto-PDIM forms were upregulated in Δ*pknF* strains. (**D**) Mass spectral intensity for C93 keto-PDIM detected in mass normalized lipid extracts from wild type Mtb (black), Δ*pknF* (red) and Δ*pknF*-C (blue) are shown in biological triplicate. (**E**) The indicated bacterial strains were grown in the presence or not of vancomycin for 7 days and their relative growth was measured via optical density at 600 nm. Statistical analysis was performed via one-way ANOVA (B) or negative binomial (E) (* p<0.05, ** p<0.01, *** p<0.001, **** p<0.0001).

Whereas TLC identification relies on interpretations of banding patterns and comigration with standards, mass spectrometry (MS) analysis can provide direct detection of molecules based on their accurate mass, which allows calculation of their chemical formulas, and, with collision-induced dissociation (CID), detection of chemical components of macromolecules for direct and unambiguous chemical assignment of the identity and substructure of molecules detected. Positive mode electrospray ionization (ESI) time of flight (TOF) MS detected an ion of *m/z* 1397.433 that matched the ammonium adduct of PDIM ([M+NH3], C93H18205). CID-MS detected neutral loss fragments corresponding to the loss of one (m/z 941.962) or two (m/z 503.519) C29 mycocerosate chains, allowing a deduced assignment of a C35 keto-modified backbone (C93 keto-PDIM) (Fig. 1B). Further confirming the match with the expected patterns of a natural Mtb-produced PDIM lipid family, we also detected a mass matching the ammonium adduct of a C93 methoxy PDIM (Fig. 1C) and its expected alkyforms. For example, we detected two alkane series, corresponding to the expected methoxy- and keto-PDIM backbones. We further detected 12 chain length analogs in two alkane series, where the median (C94) and range (C88-101) of length match the reported length of PDIM produced by Mtb when grown in culture (43, 44). Finally, the spectral intensity counts in high-performance liquid chromatography-MS analysis of a representative C93 keto-PDIM for Mtb (black), Δ*pknF* (red) and Δ*pknF*-C (blue) derived from three independent total lipid extractions showed that PDIM signals were significantly elevated in the Δ*pknF* mutant when compared to the wild-type strain, and complementation reversed the phenotype (Fig. 1D). For functional validation, we performed a vancomycin resistance assay as described (39), since PDIM levels positively correlate with vancomycin resistance (39). The Δ*pknF* strain exhibited an almost 2-fold increase in vancomycin resistance compared to wild-type or Δ*pknF*-C strains (Fig. 1E). Taken together, our results demonstrate that the deletion of *pknF* leads to a significant increase in PDIM, identifying PknF as a negative regulator of PDIM pools in Mtb CDC1551.

Given the large fold-change in PDIM pools (Fig. 1D), we hypothesized that the observed increase in PDIM of the Δ*pknF* strain would correspond to increased virulence in mice after Mtb infection, due to the known role of PDIM in promoting *in vivo* virulence (40–42). To test this hypothesis, we infected C57Bl/6 mice with 100-200 colony-forming units (CFU) (day 1 post-infection CFU, not shown) of the indicated bacterial strains via intrapharyngeal pulmonary delivery. After 28 days, the CFU in the bronchoalveolar lavage fluid (BALF) (Fig. 2A), lungs (Fig. 2B) and spleen (Fig. 2C) were 100-fold increased in mice infected with Δ*pknF* strain compared to Mtb or the complemented strain. The increase in CFU corresponded with significantly increased inflammation as assessed by hematoxylin and eosin staining of lung sections (Fig. 2D+E). To further assess changes in host responses we measured inflammatory cytokines in the lungs of mice infected with either Mtb, Δ*pknF* strain, or the complemented mutant strain (Fig. 2F-H and Supp. Fig 1). The proinflammatory cytokines MCP-1, CXCL10, GM-CSF, IFNγ and TNFα all showed a significant increase in Δ*pknF* mutant-infected mice when compared to wild-type and Δ*pknF*-C-infected lungs (Supp. Fig. 1A-E) while IL-4, IL-6, IL-10, IL12p70a and IL-17 showed no statistically significant differences (Supp. Fig. 1F-J). Similarly, in addition to IL-1β (Fig. 2F), the IL-1 family members IL-1α (Fig. 2G) and IL-33 (Fig. 2H) were both significantly elevated in the lungs of Δ*pknF*-infected mice, likely correlating with increased tissue necrosis and bacterial burden.

**Fig. 2:**
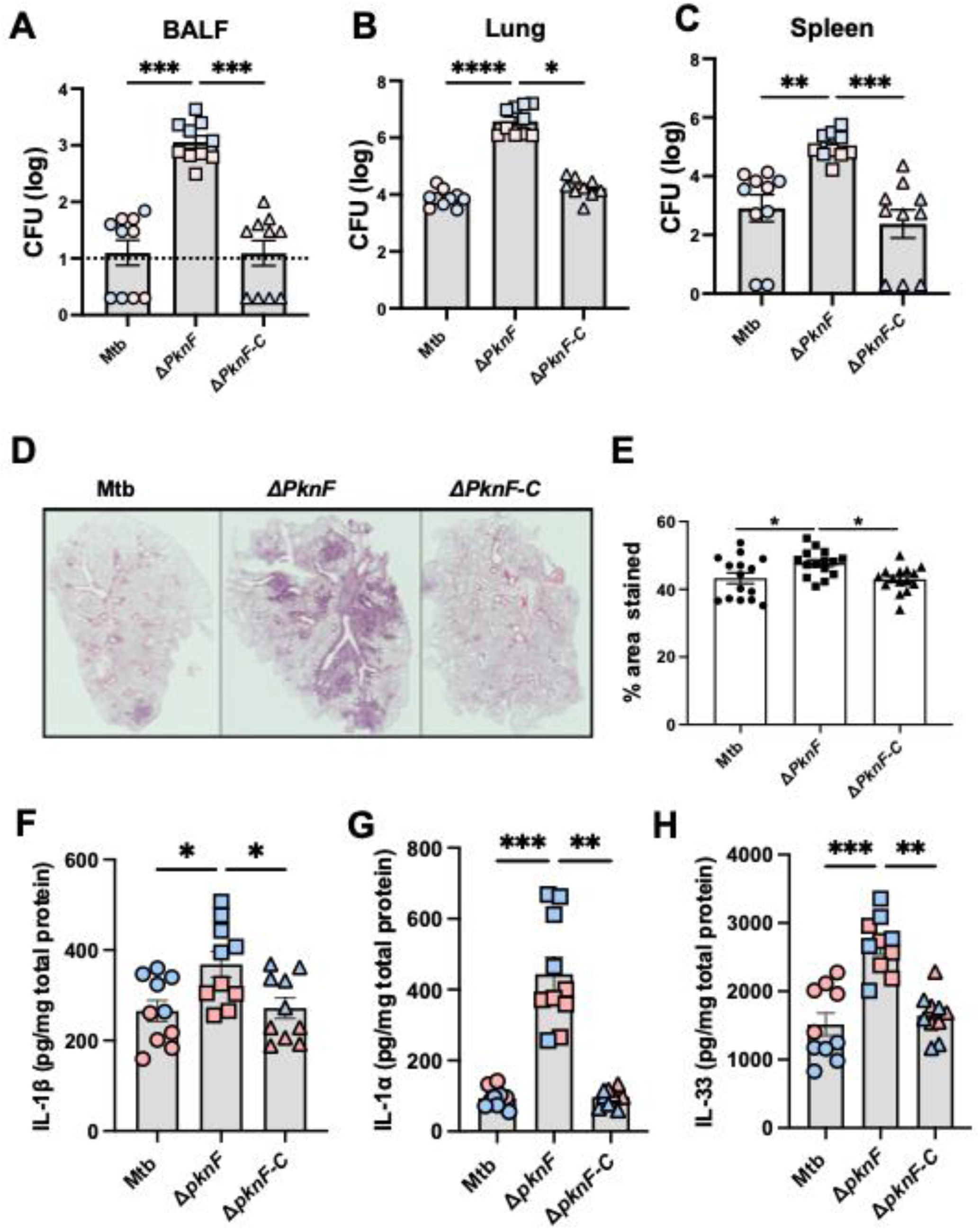
The deletion of *pknF* causes an increase in the virulence of Mtb in C57Bl/6 mice. Mice were infected with 100-200 CFU (day 1 post-infection CFU, not shown) of the indicated bacterial strains via intrapharyngeal delivery. After 28 days, the colony-forming units (CFU) in the (**A**) bronchoalveolar lavage fluid (BALF), (**B**) the lungs, and (**C**) spleen were determined. Each point represents one infected mouse from two independent infections (blue and pink colors). (**D+E**) Three slices per lung from 5 different mice were stained with hematoxylin and eosin. Image quantification was done via ImageJ. (**F-H**) Multiplexed ELISA (Luminex) was used to determine the amount of indicated cytokine in the lungs normalized to total protein. Normality of each data set was tested. Kruskal-Wallis with Dunn’s post-test (A-C) and one-way ANOVA with Tukey’s post-test (D-H) were performed to compare experimental groups (* p<0.05, ** p<0.01, *** p<0.001, **** p<0.0001).

To experimentally test whether the detected increase in bacterial burden and IL-1β in the lungs of mice infected with Δ*pknF* was due to the lack of the ability to suppress NLRP3 inflammasomes as shown before *in vitro* (1), we next infected *Nlrp3* knock-out mice (*Nlrp3*^-/-^) or C57Bl/6 mice (WT B6) with 100-200 CFU (day 1 post-infection CFU, not shown). Similar to our observations after 28 days, three months after infection, the BALF (Fig. 3A) exhibited a 100-fold increase for Δ*pknF* bacteria compared to Mtb and Δ*pknF*-C. This increase was between 10 to 50-fold in the lungs (Fig. 3B) and spleen (Fig. 3C). Importantly, deletion of *Nlrp3* did not significantly change the amount of Δ*pknF* bacteria in the BALF, lungs, or spleen (Fig. 3A-C), arguing that dysregulated NLRP3 activation alone was unlikely to cause the loss of bacterial control in mice infected with Δ*pknF* Mtb. In line with our observations above, hematoxylin and eosin staining revealed increased lung inflammation and involvement in the Δ*pknF* mutant infected mice compared to the wild-type Mtb and Δ*pknF*-C, that was unaffected by the deletion of *Nlrp3* (Fig. 3D+E). Despite increased bacterial loads at three months after infection in the lungs of mice infected with Δ*pknF* compared to Mtb and Δ*pknF*-C, we observed no changes in IL-1β levels in the lungs of WT B6 mice infected with Δ*pknF* compared to Mtb or d Δ*pknF*-C (Fig. 3F), while the alarmins IL-1α (Fig. 3G) and IL-33 (Fig. 3H) remained elevated. However, while bacterial burden was similarly increased in Δ*pknF*-infected *Nlrp3^-/-^* mice compared to Mtb and Δ*pknF*-C, IL-1β protein in the lungs of Δ*pknF*-infected *Nlrp3^-/-^* mice was significantly reduced compared to Mtb and Δ*pknF*-C-infected *Nlrp3^-/-^* mice (Fig. 3F). These results suggest that while the control of bacterial replication via PknF may occur independently of NLRP3, the modulation of IL-1β production via PknF required NLRP3 *in vivo,* as previously documented *in vitro* using bone marrow-derived macrophages (1) and dendritic cells (2). In contrast, protein levels of IL-1α (Fig. 3G) and IL-33 (Fig. 3H) were increased in ΔpknF Mtb-infected *Nlrp3^-/-^* lungs compared to lungs from *Nlrp3^-/-^* mice infected with Mtb or Δ*pknF*-C. No differences were observed in MCP-1 (Supp. Fig. 2A), CXCL10 (Supp. Fig. 2B), GM-CSF (Supp. Fig. 2C), TNFα (Supp. Fig. 2D), IFNγ (Supp. Fig. 2E), IL-17 (Supp. Fig. 2F), IL-4 (Supp. Fig. 2G), IL-6 (Supp. Fig. 2H), IL-10 (Supp. Fig. 2I) or IL-12 (Supp. Fig. 2J) between Δ*pknF*-infected WT B6 or *Nlrp3^-/-^* mice. Of note, IL-17, an IL-1 inducible cytokine, was decreased in the lungs of Δ*pknF* infected *Nlrp3^-/-^*mice compared to Mtb-infected *Nlrp3^-/-^* mice (Supp. Fig. 2F). In both WT B6 or *Nlrp3^-/-^* mice levels in IL-6 and IL-12p70 were reduced in Δ*pknF*-infected lungs compared to lungs of mice infected with Mtb or Δ*pknF*-C (Supp. Fig. 2H+J).

**Fig. 3:**
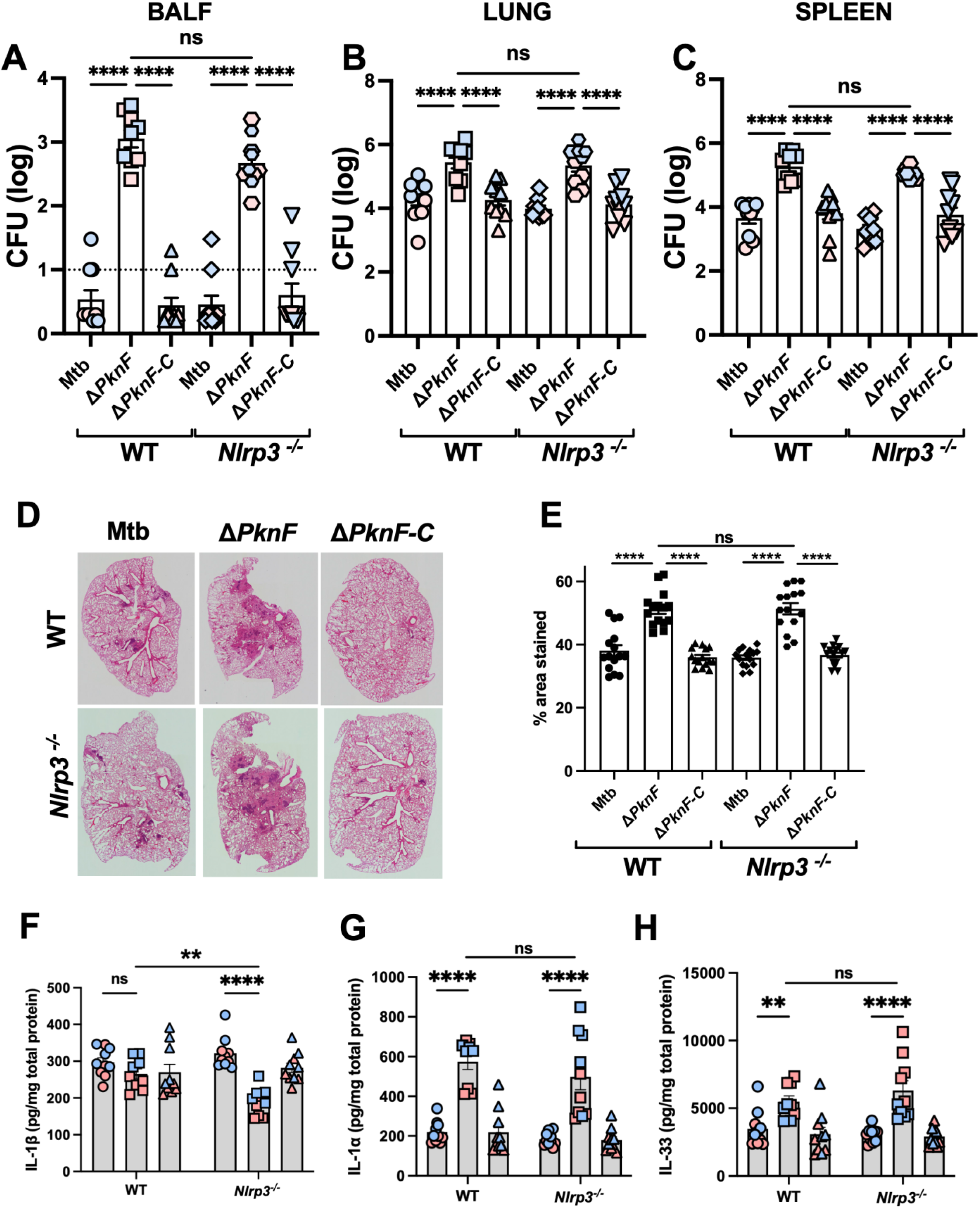
The increased virulence of the MtbΔpknF mutant is independent of NLRP3. C57Bl/6 mice (WT) and *Nlrp3* knock-out mice (*Nlrp3^-/-^*) were infected with 100-200 CFU of the indicated bacterial strains via intrapharyngeal delivery. After 90-98 days, the colony-forming units (CFU) in the (**A**) bronchoalveolar lavage fluid (BALF), the (**B**) lungs and (**C**) spleen were determined. Each point represents the result of one infected mouse from two independent infections (blue and pink colors). (**D+E**) Three slices per lung from 5 different mice were stained with hematoxylin and eosin. Image quantification was done via ImageJ. (**F-H**) Multiplexed ELISA (Luminex) was used to determine the amount of indicated cytokine in the lungs normalized to total protein. Circles indicate wild-type Mtb, squares the Δ*pknF* strain and triangles the complemented Δ*pknF* strain. Normality of each data set was tested and two-way Anova with Tukey’s post-test was performed to compare experimental groups (* p<0.05, ** p<0.01, *** p<0.001, **** p<0.0001).

Finally, to assess the virulence of the Δ*pknF* strain in the context of lower host resistance, we utilized highly susceptible mice carrying the C3HeB/FeJ-derived sst1-susceptible (sst1^S^) locus on a B6 background (B6.Sst1^S^ mice) that develop well-organized necrotic granulomas and succumb rapidly to Mtb infection (45). Consistent with the observed increase in bacterial loads in both WT B6 or *Nlrp3^-/-^* mice, host resistance in B6.Sst1^S^ mice was reduced with significantly increased moribundity in mice infected with around 60 CFU of Δ*pknF* compared to Mtb or Δ*pknF*-C (Fig. 4A). When mice were infected with a tenfold higher dose, the median time to moribundity of Δ*pknF*-infected mice was only 31 days (Fig. 4B) compared to 97 days with 60 CFU infection. The higher dose experiment was terminated after 130 days since none of the wild-type Mtb-infected mice had died by then, and only one of the Δ*pknF*-C-infected mice. These data support that Mtb PknF is a potent modulator of virulence and host resistance in both resistant and susceptible mice.

**Fig. 4:**
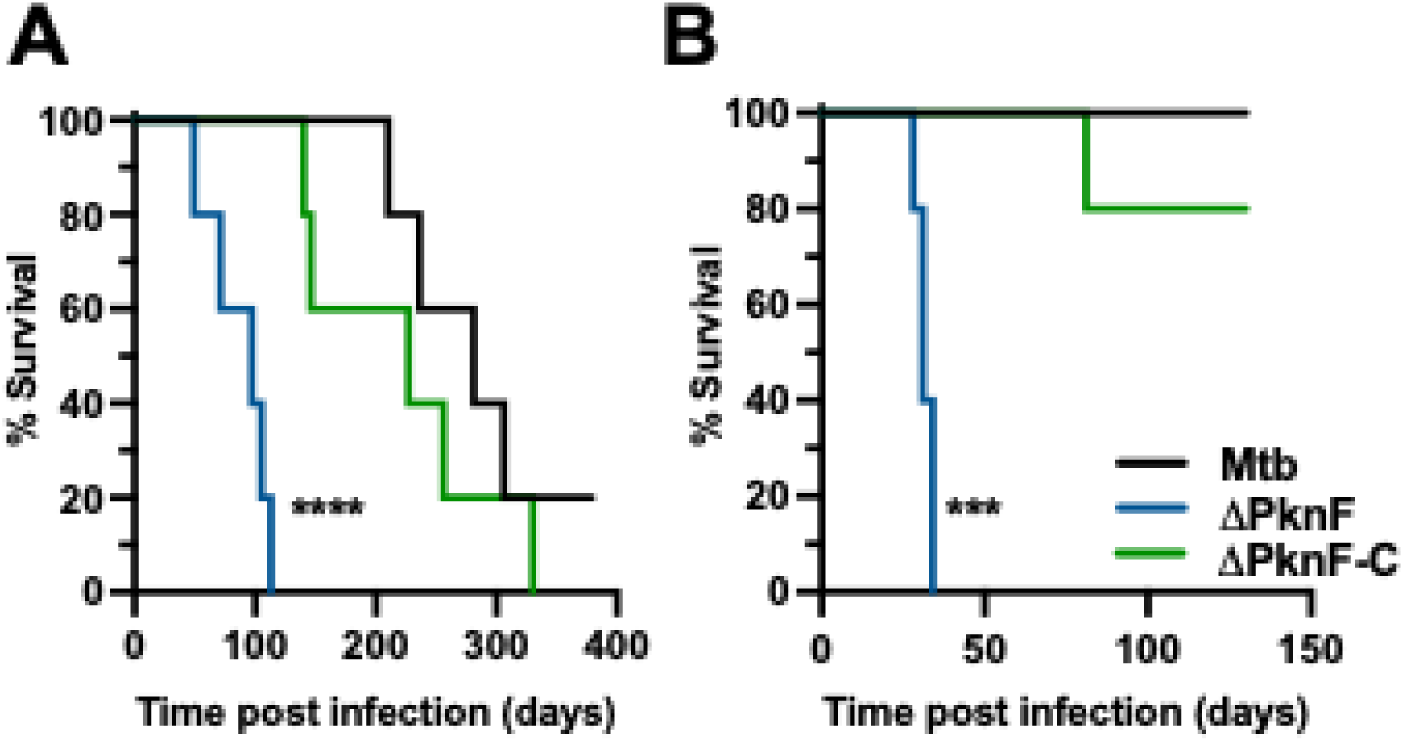
Enhanced virulence and reduced host resistance of the MtbΔ*pknF* mutant in B6.Sst1^S^ mice. Mice (n=5) were infected with (**A**) about 60 CFU or (**B**) about 600 CFU of Mtb (black), Δ*pknF* (blue) or a complemented Δ*pknF* strain (Δ*pkn*F-C, green). The survival was monitored for the indicated times. Survival curves were compared using the Log-rank Mantel-Crox test (*** p<0.001, **** p<0.0001).

## Discussion

We identified Mtb *pknF* in a screen for Mtb genes that mediate host cell inflammasome inhibition *in vitro* (1). We now show that the deletion of *pknF* significantly increases the levels of PDIM in the Mtb CDC1551 strain. Acting as a long-chain polyketide lipid that resides near the surface of Mtb, PDIM was one of the first genetically assigned virulence-controlling lipids (41, 42). PDIM loss occurs commonly during *in vitro* growth of Mtb (38). In the past this has led to potential for incorrect assignment of virulence effects to certain gene just because the Mtb deletion mutant had lost its PDIM expression. It is thus notable that in our data the complementation of *pknF* deletion reverses biochemical PDIM phenotypes as measured by TLC detection and HPLC-MS detection, as well as functional phenotypes like vancomycin response, *in vivo* immune responses, and bacterial survival.

Further, complete PDIM loss due to inactivation of any of the many genes required for its biosynthesis commonly occurs during *in vitro* growth of Mtb (38), and other work showed physiological regulation across a spectrum of biosynthesis levels (39). In this setting of an emerging understanding that this crucial virulence lipid is actively tuned by Mtb, our results suggest a specific mechanism of tuning whereby expression of the serine/threonine protein kinase PknF downregulates PDIM levels, inviting future work to identify counterregulatory activators of PDIM biosynthesis in Mtb that may either act directly through PknF or act independently of this regulatory pathway. For example, the deletion of protein kinase H (*pknH*) of Mtb results in a decrease in PDIM production (46). The precise mechanisms for this decrease remain unclear. Both PknF and PknH belong to the same family of serine/threonine protein kinases, arguing that enzymes involved in the biosynthesis or degradation of PDIM are regulated by serine/threonine phosphorylation. In fact, phosphoproteomic analysis of a *ΔpknF* Mtb strains showed a reduced phosphorylation of serine residue 447 in FadD28, an enzyme involved in the biosynthesis of PDIM (28). The phosphorylation of serine 499 of FadD28 is upregulated in the *ΔpknH* deletion mutant, suggesting an indirect effect of PknH on the phosphorylation of this site (28). The transcriptional repressor, Rv3167c, suppresses genes in the PDIM biosynthesis and transport operon, leading to increased PDIM levels in the *Δrv3167c* Mtb mutant (47). A single nucleotide polymorphism in the β subunit of RNA polymerase gene (*rpoB*), that is the most common cause of acquired rifampin resistance, leads to increased PDIM levels (48, 49) via transcriptional upregulation of genes involved in PDIM biosynthesis (49). The question remains as to how precisely these pathways that regulate PDIM levels are interconnected and potentially employed by Mtb during infections *in vivo*.

PDIM is a major virulence determinant of Mtb (40–42). We show here that the Δ*pknF* mutant Mtb strain, with its increased PDIM expression, exhibited increased virulence accompanied by decreased host resistance as demonstrated by increased pulmonary bacterial loads and significantly decreased survival in infected mice compared to the parental Mtb strain. These results are consistent with our previous finding that the Δ*rv3167c* Mtb strain, which has increased PDIM, was more virulent in the C57Bl/6 mouse model after aerosol infection (50). The *ΔpknH* strain of Mtb, with lower PDIM expression (46), was more virulent in BALB/c mice after intravenous infections (51). This discrepancy could be due to a different route of infection or that PknH affects the virulence of Mtb in other ways, independent of PDIM synthesis. Despite all these compelling data on the importance of PDIM for virulence in the mouse model, few data are available on the molecular mechanisms underlying this phenotype *in vivo*. There are numerous studies assigning multiple functions to PDIM during Mtb’s *in vitro* interaction with macrophages (52). One of these is its capacity to limit autophagy in human macrophages (53). A genetic screen revealed that PDIM is also important for limiting autophagy-mediated killing of Mtb in murine BMDMs (54). Importantly, this study clearly delineated the importance of PDIM-mediated autophagy inhibition for the *in vivo* virulence of Mtb (54).

Another prominent PDIM function is its role in permeabilizing the phagosomal membrane and allowing for escape of the bacteria from the phagosome (47, 55–58). The escape of Mtb from the phagosome has been correlated with its capacity to induce cell death (47, 50, 59, 60). The high PDIM level Mtb mutant Δ*rv3167c* increased host cell necrosis via an uncharacterized cell death pathway (50). We also observed an increase in cell death in the lungs of mice infected with the Δ*pknF* Mtb mutant. We previously demonstrated that in BMDMs and BMDCs the Δ*pknF* mutant induces an increase in pyroptosis by activating the NLRP3-inflammasome (1, 2). Our data suggest that *in vivo*, pyroptosis is not a major cell death pathway induced by Δ*pknF*, since the increased virulence and bacterial burdens of the mutant are similar between WT B6 and *Nlrp3^-/-^* mice. Nevertheless, and consistent with our initial *in vitro* observations (1), Nlrp3 inflammasome-dependent IL-1β production remained PknF dependent in the lungs of mice, since *Nlrp3^-/-^* mice infected with Δ*pknF* had significantly lower levels of IL-1β in the lungs than *Nlrp3^-/-^* mice infected with Mtb or Δ*pknF*-C.

Overall, our data reveal a previously unrecognized role for PknF in the modulation of bacterial virulence *in vivo.* In addition, our results reveal new directions for future mechanistic studies examining the role of PknF in Mtb lipid modulation that might focus on the regulation of PDIM production by PknF kinase-mediated action on other gene products.

## Materials and Methods

### Mice

C57BL/6NTAC mice were obtained from Taconic Bioscience (NY, USA). C57BL/6 mice expressing a Foxp3-GFP reporter (C57BL/6-Foxp3^tm1Kuch^ (61) or Thy1.1 (B6.PL-*Thy1^a^*/CyJ) were also used as wild type (WT B6) C57BL/6 controls with similar results to C57BL/6NTAC mice. C57BL/6-Foxp3^tm1Kuch^, B6.PL-*Thy1^a^*/CyJ, *Nlrp3^-/-^* (B6N.129-*Nlrp3^tm2Hhf^*/J,{ 2009.Brydges}, and B6 Sst1 (B6J.C3-sst1^C3HeB/Fej^ (45), were obtained through a supply contract between NIAID and Taconic Farms. Mice of both sexes were used and 12-18 weeks old at the onset of experiments. Mice within experiments were age and sex matched.

### Ethics Statement

All animals were bred and maintained in an AAALAC-accredited ABSL2 or ABSL3 facility at NIAID and experiments were performed in compliance with an animal study proposal approved by the NIAID Animal Care and Use Committee.

### Mtb infection of mice

Mtb CDC1551 WT, *pknF* deletion mutant (MtbΔ*pknF*) and *pknF* complemented (Δ*pknF*-C) strains (1) were grown in Middlebrook 7H9 broth (Sigma Aldrich, Darmstadt, Germany) supplemented with 10% OADC, 0.5% of glycerol and 0.05% Tween-80; for mutant and complement strains, the antibiotics hygromycin (50µg/mL) and kanamycin (50µg/mL) were added, respectively. Bacterial concentration in log phase was determined by OD_600_. Briefly, once bacteria reached OD=0.6, bacteria were centrifuged at 5000rpm for 10min. Then, resuspended in PBS and flushed through a needle 27g. Finally, the bacteria were diluted in PBS targeting 100-200 of colony forming units (CFU) per 50µl of inoculum per mouse. The bacteria were delivered to the lungs of mice via intrapharyngeal (i.ph.) route, and infection doses were confirmed by plating lung homogenates one day post-infection.

### CFU determination

Lung and spleen homogenates were generated using Precellys Evolution machine (Bertin Instruments, France). Serial dilutions of homogenized tissue and bronchoalveolar lavage fluid (BALF) were done in PBS-0.05% Tween-20, then plated on Middlebrook 7H11 agar plates (Sigma Aldrich, Darmstadt, Germany) supplemented with 10% OADC (Oleic Acid-Albumin-Dextrose-Catalase; Beckton Dickenson, MD, USA) and 0.5% glycerol (Invitrogen, Karnataka, India). CFU were counted after 21 days of incubation at 37°C.

### Lung Cytokine/Chemokine Quantification and Histology

Lung homogenates were generated using Precellys Evolution machine (Bertin Instruments, France). Samples were centrifuged and sterile-filtered using 0.22µm filter and then stored at −80°C. The cytokine/chemokine analysis was done by using the Luminex MAGPIX® platform according to manufacturer protocol (R&D Bioscience). BCA assays were performed on lung homogenates to normalize cytokine levels as pg per mg of total protein.

The right lower lobes of the lungs were fixed in 10% buffered formalin for histopathology. Lung lobes were washed and suspended in PBS, followed by paraffin embedding, sectioning and hematoxylin and eosin (H&E) staining done by Histoserv Inc (https://www.histoservinc.com/). The tile scanning and image stitching were performed for each mouse lung section, and slides were imaged on Nikon Eclipse microscope. The images were generated using Nikon’s NIS-Elements imaging software. Quantification of H&E-stained lung sections was done using Image J software.Paraffin embedding, sectioning and hematoxylin and eosin staining were done by Histoserve Inc (https://www.histoservinc.com/).

### Lipid Analysis

Lipid extracts were created as described (43) and analyzed using an Agilent 6546 Q-Tof mass spectrometer connected to an Agilent Infinity II liquid chromatography system using normal-phase chromatography was performed as described (43). Total lipid extracts of WT Mtb, ΔpknF and ΔpknF-C strains were analyzed for PDIM levels using one-dimensional TLC. PDIMs were developed using petroleum ether:diethyl ether (9:1) and lipid spots were visualized by staining with 5% solution of molybdophosphoric acid in 95% ethanol followed by charring at 140°C Reversed phase chromatography was done on an LCMS system using an Agilent Poroshell 120 column (EC-C18, 1.9 mm 3 x 50 mm) with gradient elution. Mobile phase A was 5:95 water:methanol and mobile phase B was 85:15 n-propanol:cyclohexane. The gradient started at 100% A, from 2 min to 15 min it increased to 100%B, holding at 100% until 23 min. It then returned to 100% A and held from 25-30 min to recondition the column. The flow rate was 0.15 mL/min throughout the run. Collisionally induced dissociation was done using a narrow (1.3 *m/z*) isolation window and 25 V collision energy.

To assess PDIM levels via a functional assay, we performed a vancomycin resistance assay as described (39).

### Statistical analyses

Statistical analysis was performed using GraphPad Prism version 10.0 software. Data was analyzed using Kruskal-Wallis with Dunn’s post test. One-way or two-way ANOVA with Tukey post-test was performed as indicated after normal distribution was confirmed using Shapiro-Wilk test and QQ plot analysis. Time to moribundity studies were assessed using the Log-rank Mantel-Cox test for survival. p-value significance is as follows - *- ≤0.05, ** - ≤0.01, *** - ≤0.001, **** - 0.0001.

## Acknowledgments

Research was supported by NIH/NIAID grant R01AI147630 (VB, SR, VKB, and SEK) and R01 AI165573 and U19 AI162584 (DBM). The funders had no role in study design, data collection, and interpretation, or the decision to submit the work for publication. This research was supported in part by the Intramural Research Program of the National Institutes of Health (NIH). The contributions of the NIH authors (FTJ, PJB, KDMB) are considered Works of the United States Government. The findings and conclusions presented in this paper are those of the authors and do not necessarily reflect the views of the NIH or the U.S. Department of Health and Human Services.

---

**Supp. Fig. 1:**
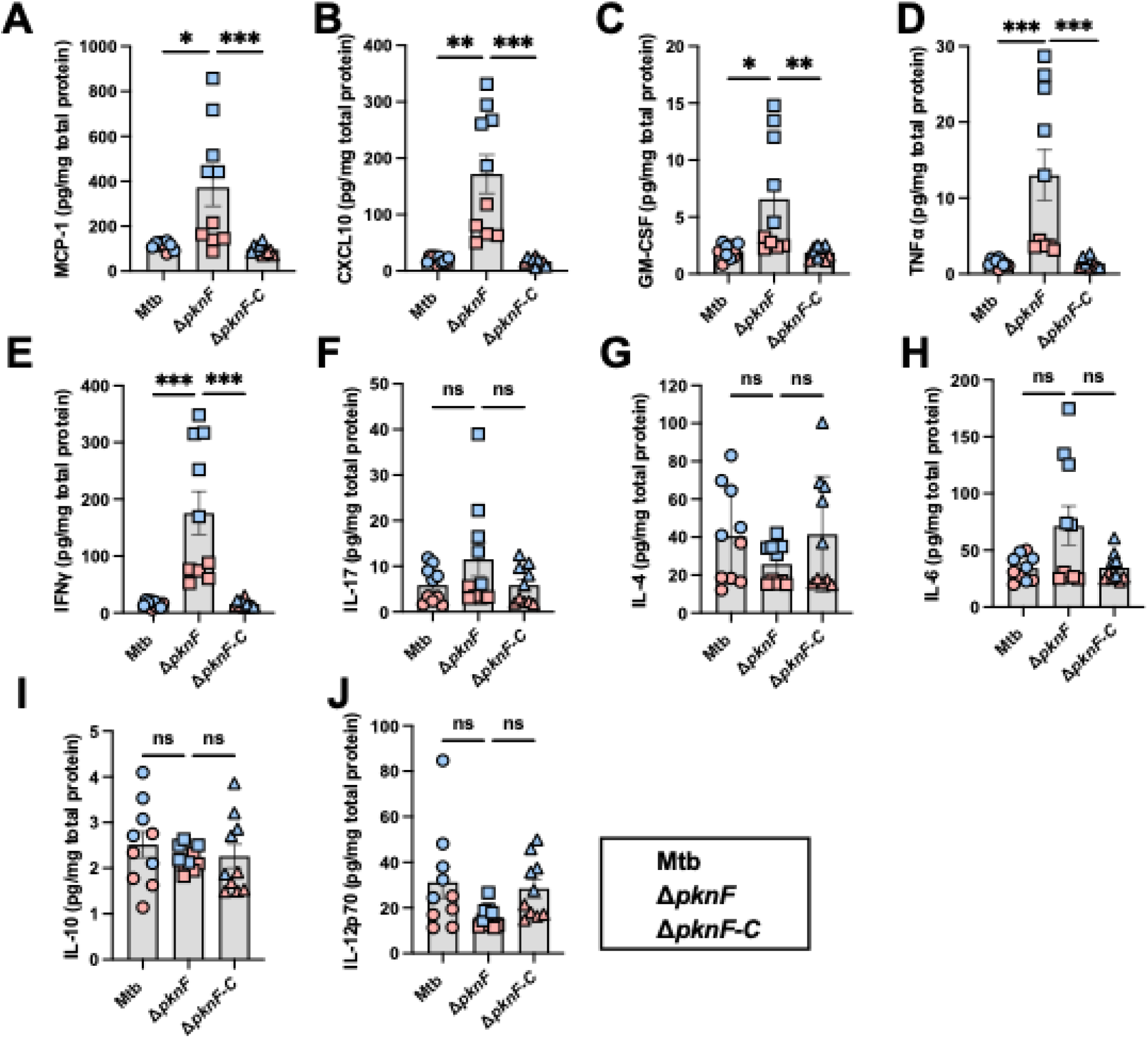
Characterization of cytokine levels in lungs at day 28 after Mtb infection. Multiplexed ELISA (Luminex) was used to determine the amount of indicated cytokine in the lungs normalized to total protein. Each point represents one infected mouse from two independent infections (blue and pink colors). Normality of each data set was tested. (A-E, G, H) Kruskal-Wallis with Dunn’s post-test and (F, I, J) one-way ANOVA with Tukey’s post-test were performed to compare experimental groups (* p<0.05, ** p<0.01, *** p<0.001, **** p<0.0001).

**Supp. Fig. 2:**
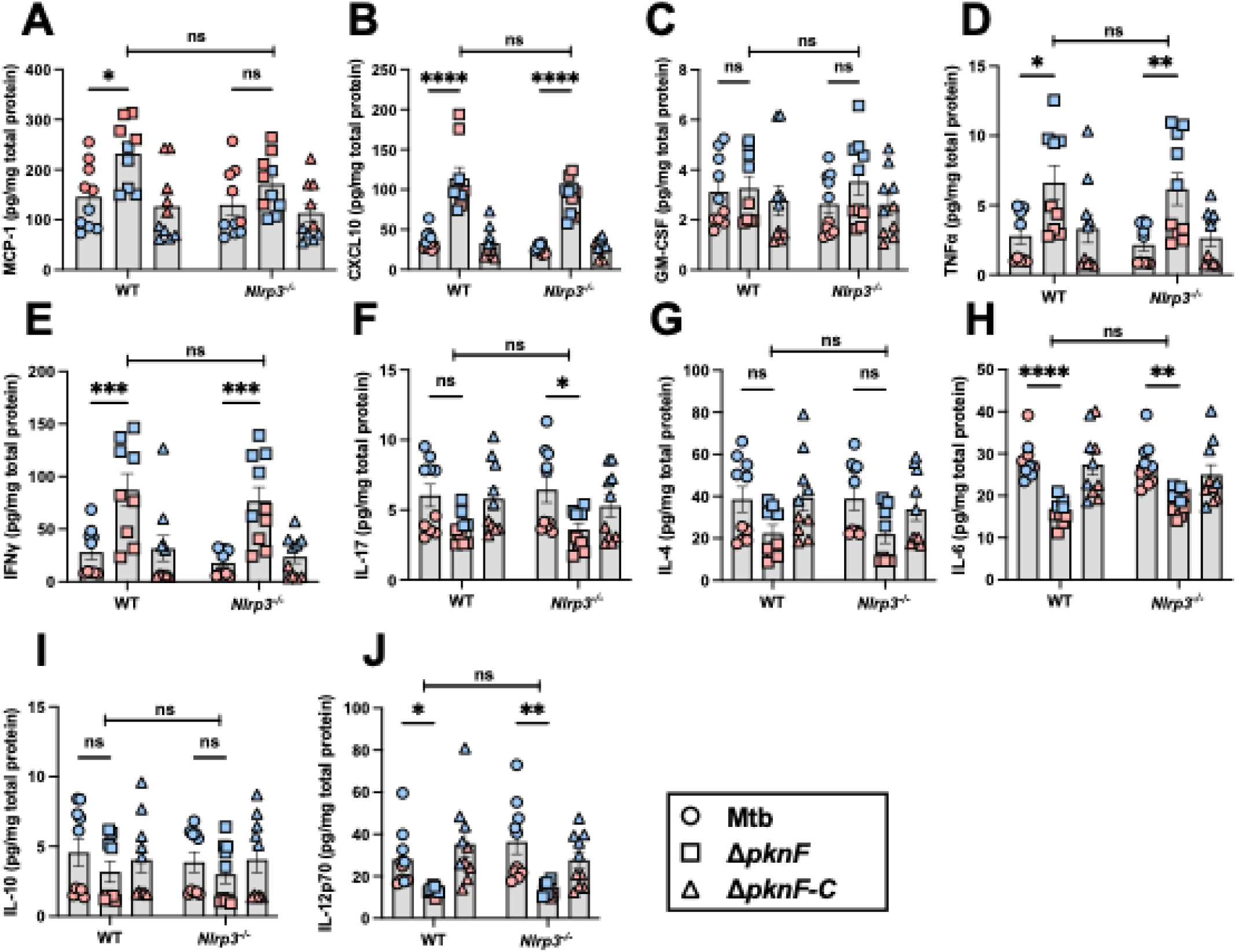
Characterization of cytokine levels in lungs of wild-type and *Nlrp3^-/-^*mice at day 90-98 after Mtb infection. Multiplexed ELISA (Luminex) was used to determine the amount of indicated cytokine normalized to total protein in the infected lungs described in Figure 3. Each point represents the result of one infected mouse from two independent infections (blue and pink colors). Normality of each data set was tested, and two-way ANOVA with Tukey’s post-test was performed to compare experimental groups (* p<0.05, ** p<0.01, *** p<0.001, **** p<0.0001).

